# Antibacterial activity of *Balanites aegyptiaca* Oil Extract on *Staphylococcus aureus* and *Escherichia coli*

**DOI:** 10.1101/2021.03.23.436600

**Authors:** Sabina Khanam, Fatima Zakari’yau Galadima

## Abstract

Africa is very rich with biodiversity resources, vegetation and it is estimated about 40,000- 45,000 species of different flora. A very large number of plant species are medicinally used for the treatment of various diseases. *Balanites aegyptiaca* also called “desert date” indeed a plant with amazing benefits for human, *Balanites aegyptiaca* was popularly known to have added value and hope for person who went to pursue it for health. In Nigeria, there are evidences of serious bacterial infections with Gram-positive and Gram-negative *Staphylococcus aureus* and *Escherichia coli* respectively. *Balanites aegyptica* is used to treat so many illnesses including laxative, diarrhea, hemorrhoid, stomach aches, jaundice, yellow fever, syphilis and epilepsy. The *Balanite aegyptica* sample was sun dried for easy removal of the seed from the shell; it was washed to remove un-wanted particles and was dried off then pounded fine powder using mortar and pestle. **A**ntibacterial activity of the *Balanites aegyptica* oil at different concentrations on *staphylococcus aureus* and *Escherichia coli* bacteria and different meters in diameter of the zones was observed.The results shows significant effect of Balanites oil on bacteria by observing the presence of clear spaces known as the zone of inhibitions in the experiments.

## INTRODUCTION

Microorganisms such as bacteria, fungi, viruses and parasites are mostly pathogenic and have great impact on health because of high growing incidences of infectious diseases. In Nigeria, there are evidences of serious bacterial infections with Gram-positive and Gram-negative *Staphylococcus aureus* and *Escherichia coli* respectively (Brent *et al*., 2006; Bello *et al*., 2017; Chaturvedi *et al.*, 2009; Church and Maitland, 2014; Evans *et al*., 2004; Gwer *et al*., 2007; Uneke, 2008).

*Balanites aegyptica* seed were produced annually in the tropical and sub tropical countries of Asia and African. Like rest of the plant, they were highly valued. The *Balanites aegyptica* also known as the tree of life has a host of nutrient uses for both people and livestock a like Balanites seed have a unique and pleasing appearance. When firmly rooted, they produce a bountiful crop of one of the worlds of healthy plants.The trees itself was rather slender with dropping branches that grow to approximately 1O-12m in height. *Balanites aegyptiaca* is one of the most common but neglected wild plant species of the dry land areas of Africa and South Asia.Uses of medicinal plants for primary health care have increased in recent years worldwide. Various scientists are searching new phytochemical for the treatment of infectious diseases. Plants have wide variety of secondary metabolites that have antimicrobial properties. *Balanites aegyptica* is used to treat so many illnesses including laxative, diarrhea, hemorrhoid, stomach aches, jaundice, yellow fever, syphilis and epilepsy. It is used to treat liver disease and as a purgative and sucked by school children as a confectionary in some countries even in Nigeria The bark is used in the treatment of syphilis, round worm infection and as a fish poison. The aqueous leaf extract and spooning isolated from it kernel cakes have antibacterial activity (Doughari *et al*., 2007).

Africa is very rich with biodiversity resources, vegetation and it is estimated about 40,000- 45,000 species of different flora. A very large number of plant species are medicinally used for the treatment of various diseases. Due to easy accessibility, cheaper than synthetic drugs, less toxic, and less side effects when compared to other synthetic medicine the use of medicinal plants has increased now a days (Croft and Yardley, 2002). *Balanites aegyptica* is a highly valued plant distributed in many countries of the tropic and sub tropic, desert, etc. It has impressive range of medicinal uses with high nutritional value. Different parts of the plant contain a profile of important minerals, and a good source of protein, vitamins, amino acids and various phenols. The Balanites plant provides a rich and rare combination of zeatin, quercetin, kaempferom and many other phytochemicals.

## MATERIALS AND METHODS

### STUDY AREA

Damaturu metropolis was considered as the study area. It has an area of 737km and population of 302, 145 according to 2006 population census, it’s located in North-Eastern province of Nigeria. Damaturu is located between the latitude of 11°57’58.29’’E or 11,7746996 respectively.The elevation of Damaturu is 376.23 meters above sea level and hence the population density of 48,014 person ( census 2006). The region is a sahel Savannah which has some sustainable of the indigenous plant species.

### MATERIALS USED

1. Mueller Hinton agar
2. Autoclave
3. Hot air oven
4. Mortar and pestle
5. Conical flask
6. Cotton wool
7. Wire loop
8. Test tube
9. Weighing balance
10. Measuring cylinder
11. Aluminum foil
12. Petri-dishes
13. Incubator
14. Pipette
15. Soxhlet extractor (Apparatus)
16. Separating funnel
17. Cork borrer
18. Spirit lamp
19. Beaker
20. Sensitivity disc
21. Filter paper

### REAGENTS

1. Distilled water
2. Petroleum ether
3. Dimethyl sulfoxide
4. Oil extract

### SAMPLES COLLECTION

The sample were collected within Damaturu local government area of Yobe state in a polythene bag and brought to the laboratory for analysis.

### SAMPLE PREPARATION

The *Balanite aegyptica* sample was sun dried for easy removal of the seed from the shell; it was washed to remove un-wanted particles and was dried off then pounded fine powder using mortar and pestle.

### SOXHLET EXTRACTOR

Extraction was carried out according to the method adopted by prince (2000).

1. 125g of the crushed seed was weighted and then poured into a large size filter paper and steppled properly.
2. The filter paper containing the crushed seed was inserted into the extraction chamber of a SOXHLET extractor.
3. 250ml of petroleum ether was poured into the round button flask of the extractor.
4. The heating mantle was switched on at 40°c and allowed to heat the solvent petroleum ether
5. The oil flows through the tube of the extraction chamber into the boiling flask and the process continues.
6. The resulting mixture in the boiling flask was filtered and poured into a beaker. Then it was inserted into hot air oven at 30°c for the remaining solvent in the oil to evaporate.

### SEPARATING METHOD

After the sample was removed out of the hot air oven,it was allowed to cool. Then it was poured into a separating funnel and allowed to settle completely. Different layers were observed,the button was opened and allowed to drop off, when the residue layer was finished it was discarded and the oil was collected and ready for use.

### PREPARATION OF CULTURE MEDIA

1. 9.5g of Mueller Hinton agar was weighed and was dissolved in 250g of distilled water in a conical flask.
2. The cornical flask was closed with a cotton and foil paper then sealed with a masking tape.
3. The conical flask was placed into an autoclave and sterilised at 121°c for 15mins.
4. The autoclave was turned off and the conical flask was removed and allowed to cool for sometimes.
5. An amount of the prepared media was aseptically poured into four petridishes covered and allowed to solidify.

### PREPARATIONS OF EXTRACT STOCK SAMPLE

1. 1ml of extract was measured using a sterile pippete and poured into a clean and sterile test tube.
2. 9ml of dimethyl sulfoxide (Dmso) was measured and added on the extract test-tube, shaked and was labelled as “stock sample” test-tube at 100,000
3. Another five test-tubes were arranged in a row and labelled as test-tube 1,2,3,4 and 5.
4. The 1st test-tube:- 0.1ml of extract sample was measured and poured into the first test-tube and 0.9ml of Dmso was measured and added on it, which made it up to 1ml (100g/ml)
5. The 2nd test-tube:- using a sterile pippete, 0.2ml of extract sample was poured into the 2nd test-tube and 0.8ml Dmso was added on it, making 1ml sample (200g/ml)
6. The 3rd test-tube:- 0.3ml of extract sample was poured into the 3rd test-tube and 0.7ml of Dmso was added on it, also making 1ml sample (300g/ml)
7. The above procedure was also done for test-tube where 4ml and 5ml of sample and 6ml and 5ml of Dmso was diluted respectively(which made up to 400g/ml and 500g/ml

### INNOCULATION OF BACTERIA

1. The two petri dishes containing the solidified Mueller Hinton agar were arranged and labelled according to the species of bacterias used i.e *Escherichia coli* and *Staphylococcus aureus.*
2. One Petri dish labelled as *E.coli* and one as *Staphylococcus aureus*

1. Using strict plating method
2. Using a sterilised wire loop small amount of e.coli was inoculated in an aseptic condition into the two Petri dishes and was strictly rubbed on the surface of the media and the petridishes were closed.
3. After sterilising the wire loop again, calories of staphylococcus aureus was also inoculated through out the surface of the 2 plate and was closed.

## EXPERIMENTAL DESIGN

After inoculating, well diffusion method was used for the experiments.

### Well diffusion method

Cork borrer was sterilised and well was dugged on the two media plates (1 *E.coli* plate and 1 *Staphylococcus aureus* plate). Little amount of the diluent extract of the five test-tubes were poured into each hole for both plate which were labelled as 100mg/ml, 200mg/ml, 300mg/ml, 400mg/ml, and 500mg/ml respectively, the plates were incubated.

The above experimental procedures were repeated 3 times and the mean of each experiment was recorded.

## RESULTS AND DISCUSSION

Antibacterial activity of the *Balanites aegyptica* oil at different concentrations on staphylococcus aureus and *Escherichia coli* bacteria and different meters in diameter of the zones was observed. In some cases the higher the concentration higher the zone of inhibition (Table 5 to 8), also showed the inhibition zone sizes on the culture plates. In *Staphylococcus aureus* plate, The highest zone of inhibition occurred in experiment 2 where 6mm/dm,8mm/dm, 9mm/dm, 7mm/dm and 8mm/dm was recorded for 100mg/ml, 200mg/ml, 300mg/ml, 400mg/ml and 500mg/ml concentration respectively (Table 1 to 4). Also in *Escherichia coli*, the highest zone of inhibition occurs in experiment 2, where 7mm/dm, 11mm/dm, 9mm/dm, 5mm/dm and 4mm/dm was observed and recorded for 100mg/ml, 200mg/ml, 300mg/ml, 400mg/ml and 500mg/ml concentration respectively.

**TABLE-1.** Antibacterial activity of *Balanites aegyptica* seed oil extract against *Staphylococcus aureus* using well diffusion method:

| Concentration of extract | Zone of inhibition |
| --- | --- |
| 100mg/ml | 7mm/dm |
| 200mg/ml | 4mm/dm |
| 300mg/ml | 5mm/dm |
| 400mg/ml | 7mm/dm |
| 500mg/ml | 4mm/dm. |

**TABLE-2.** Antibacterial activity of *Balanites aegyptica* seed oil extract against *Staphylococcus aureus* using well diffusion method:

| Concentration of extract | Zone of inhibition |
| --- | --- |
| 100mg/ml | 6mm/dm |
| 200mg/ml | 5mm/dm |
| 300mg/ml | No visible zone |
| 400mg/ml | No visible zone |
| 500mg/ml | 4mm/dm |

**TABLE-3.** Antibacterial activity of *Balanites aegyptica* seed oil extract against *Staphylococcus aureus* using well diffusion method:

| **Concentration of extract** | **Zone of inhibition** |
| --- | --- |
| 100mg/ml | 6mm/dm |
| 200mg/ml | 8mm/dm |
| 300mg/ml | 9mm/dm |
| 400mg/ml | 7mm/dm |
| 500mg/ml | 8mm/dm |

**TABLE 4:** Mean±SD of Antibacterial activity of *Balanites aegyptica* seed oil extract against *Staphylococcus aureus* using well diffusion method.

| Concentration of extract | Zone of inhibition (Mean±SD) |
| --- | --- |
| 100mg/ml | 6.33±0.47 |
| 200mg/ml | 5.66±1.69 |
| 300mg/ml | 7.0±2.0 |
| 400mg/ml | 7.0±0.0 |
| 500mg/ml | 5.33±1.88 |

**TABLE-5.** Antibacterial activity of *Balanites aegyptica* seed oil extract against *Escherichia coli* using well diffusion method:

| Concentration of extract | Zone of inbition |
| --- | --- |
| 100mg/ml | 3mm/dm |
| 200mg/ml | 4mm/dm |
| 300mg/ml | 5mm/dm |
| 400mg/ml | 7mm/dm |
| 500mg/ml | 5mm/dm |

**TABLE-6.** Antibacterial activity of *Balanites aegyptica* seed oil extract against *Escherichia coli* using well diffusion method:

| **Concentration of extract** | **Zone of inhibition** |
| --- | --- |
| 100mg/ml | 7mm/dm |
| 200mg/ml | 11mm/dm |
| 300mg/ml | 9mm/dm |
| 400mg/ml | 5mm/dm |
| 500mg/ml | 4mm/dm |

**TABLE-7.** Antibacterial activity of *Balanites aegyptica* seed oil extract against *Escherichia coli* using well diffusion method:

| **Concentration of extract** | **Zone of inhibition** |
| --- | --- |
| 100mg/ml | 1mm/dm |
| 200mg/ml | 2mm/dm |
| 300mg/ml | 2mm/dm |
| 400mg/ml | 3mm/dm |
| 500mg/ml | 1mm/dm. |

**TABLE 8:** Mean±SD of Antibacterial activity of *Balanites aegyptica* seed oil extract against *Escherichia coli* using well diffusion method.

| Concentration of extract | Zone of inhibition (Mean±SD) |
| --- | --- |
| 100mg/ml | 3.66±2.49 |
| 200mg/ml | 5.66±3.85 |
| 300mg/ml | 5.33±2.86 |
| 400mg/ml | 5.0±1.63 |
| 500mg/ml | 3.33±1.69 |

The results obtained shows that *Balanites aegyptica* has an antibacterial agent that acted on bacteria.

As a general rule, plant seed extract concentrations were considered active against bacteria when the zone of inhibition was greater than 6mm.After the innoculation and incubation of the cultured plates, the observation was done after 24hours, and it was found out that there were clear spaces around the cup borrer well also known as zone of inhibition whereas the other part of the plate there were presence of bacteria growth, this indicates the effect of the oil on the bacteria. The inhibition zone was supposed to increase gradually with the increase in concentration. Although, in my experiment it was not like that, the maximum inhibition zone was observed in experiment of *Staphylococcus aureus* while in *E. coli* in experiment 2 also.

The result of the *Balanites* oil experiment were in close agreement with other finding obtained by other workers. *Indigofera* belong to family *Leguminosae* is renewed for its antimicrobial activity. By cup plate method extract of *Indigofera* was tested against Gram positive and Gram negative bacteria. Aqueous and hexane extract of *Indigofera* has the best inhibition against bacterial strains. By disc diffusion method antimicrobial activity of both aqueous and ethanol parts of the leaves of *Balanites aegyptica* against *Salmonella typhi.* The highest antimicrobial activity was noticed with ethanol extracts of *Balanites aegyptica* While aqueous extract showed low activity at 100 mg/ml dose. Various studied revealed antibacterial activity of *Balanites aegyptica* against most strains of bacteria (Daya *et al*., 2011; Doughari *et al*., 2007; Natarajan *et al*., 2010; Noor Jahan *et al*., 2012; Karuppusamy *et al*., 2002).

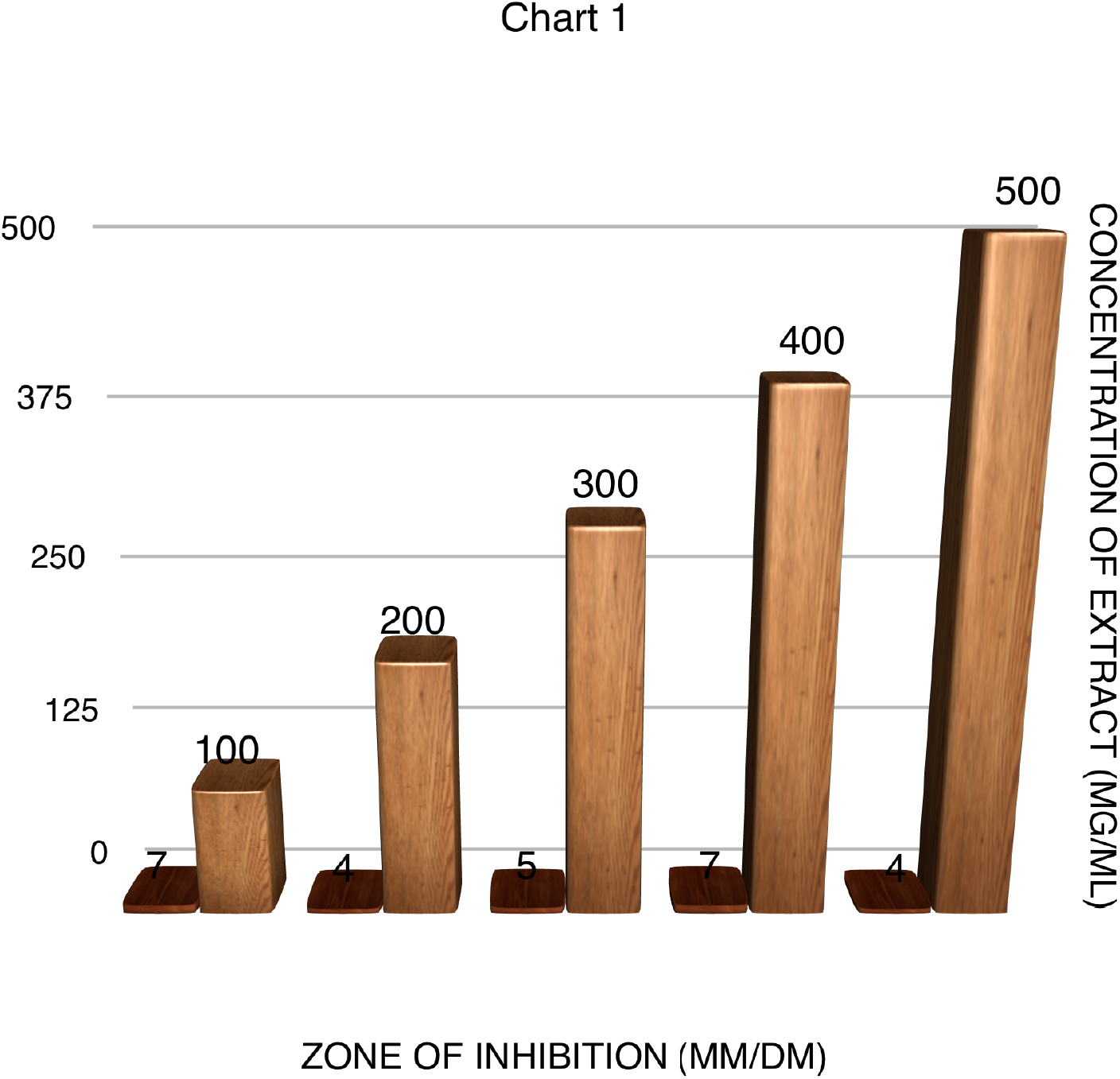

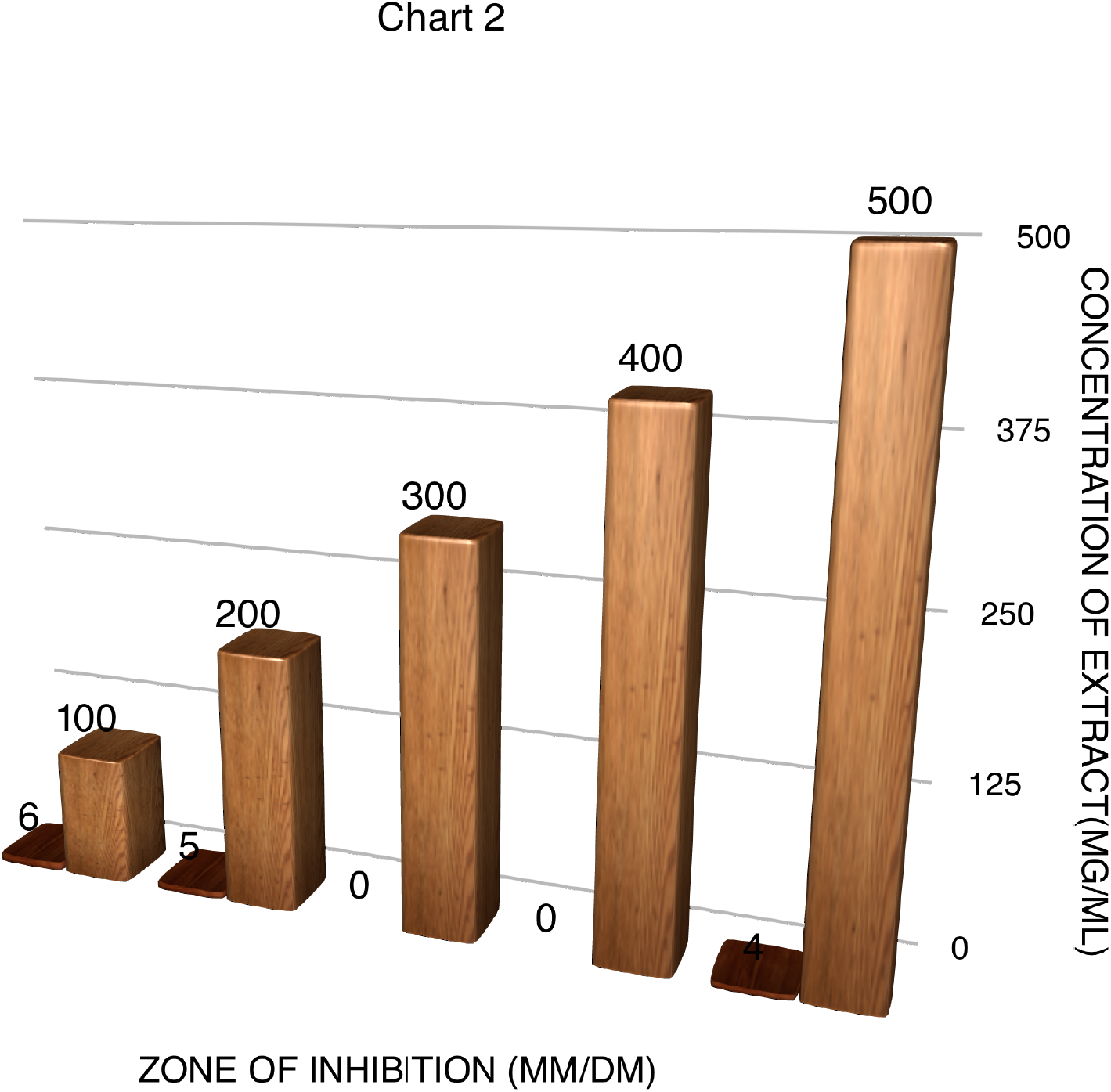

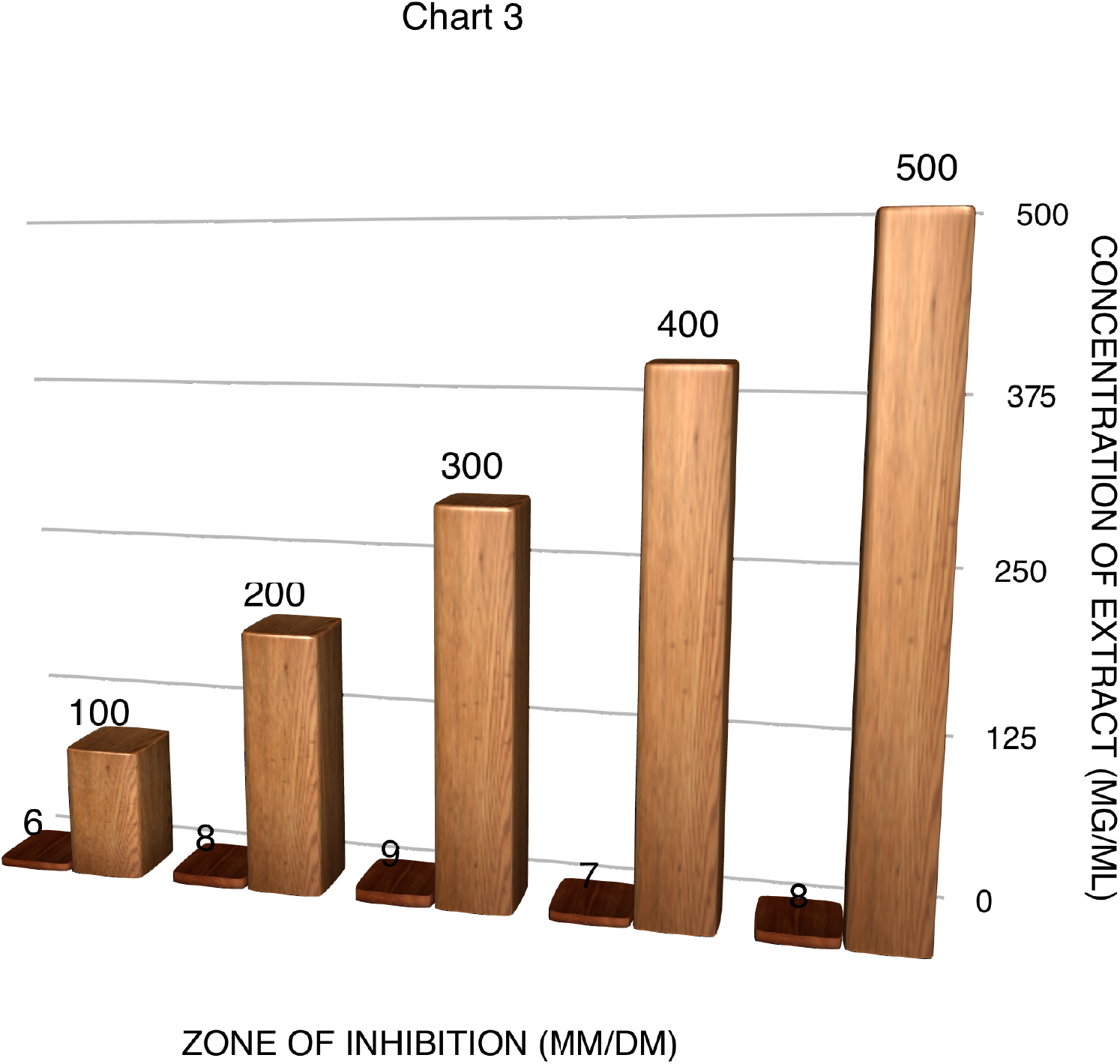

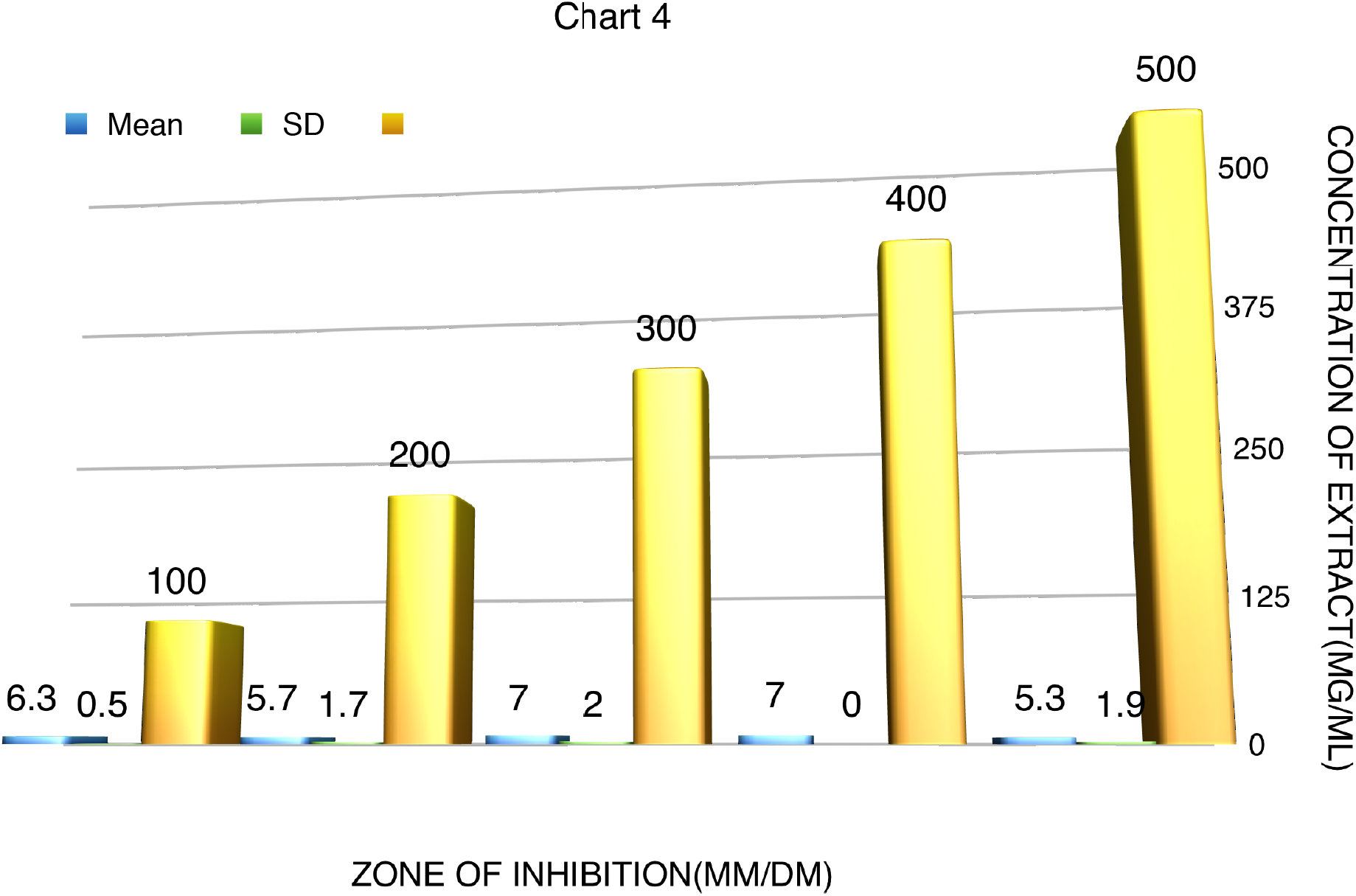

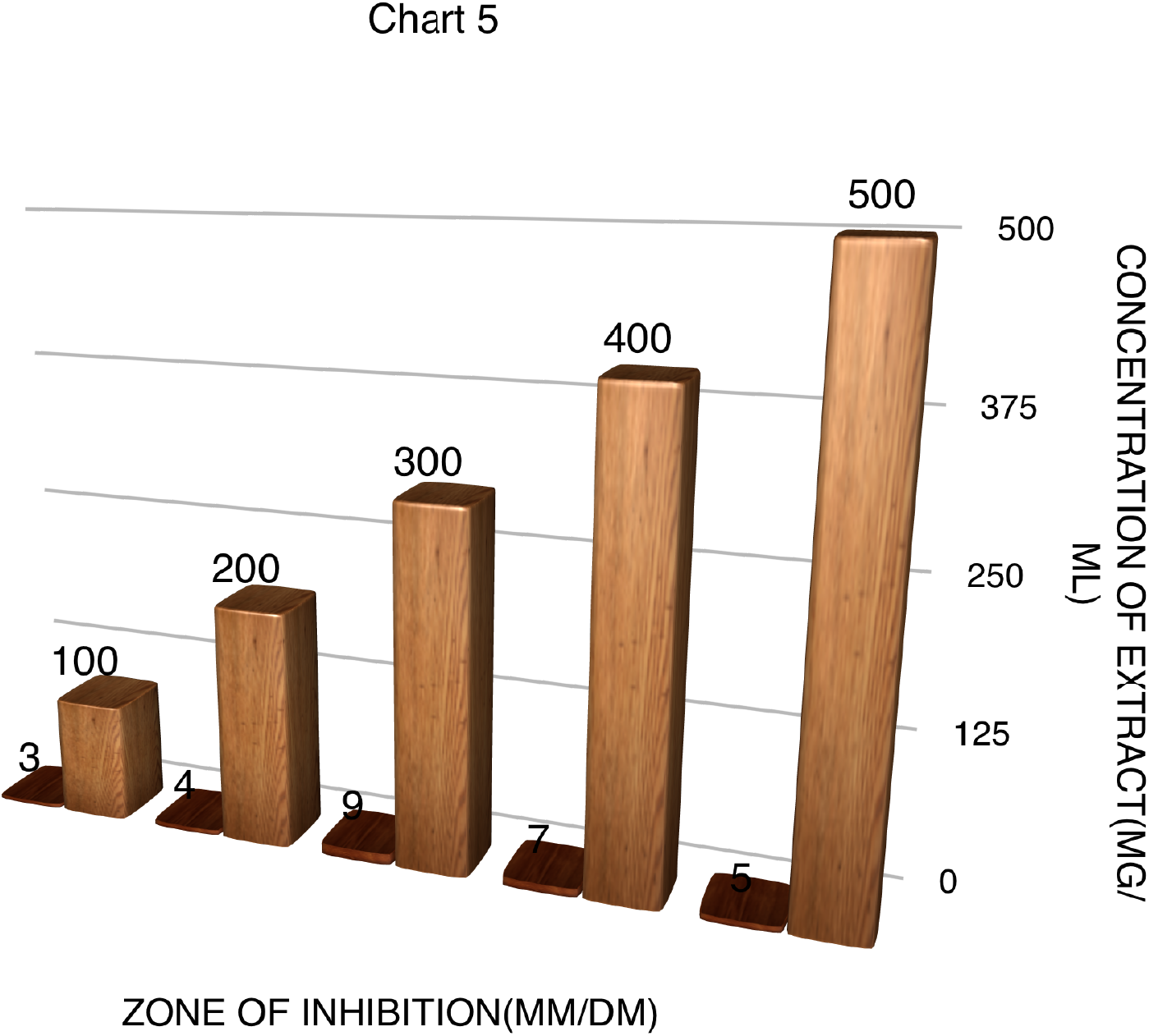

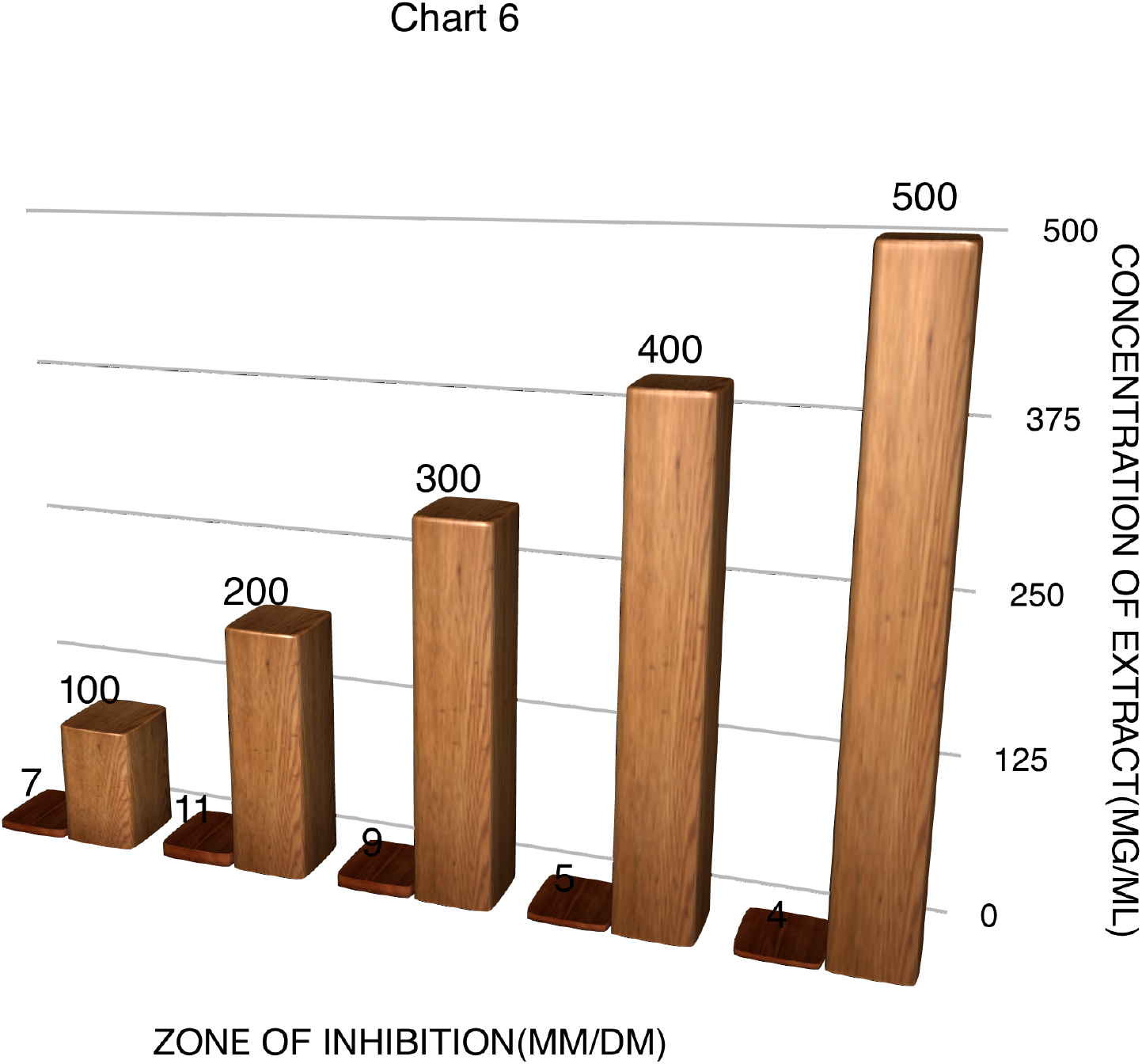

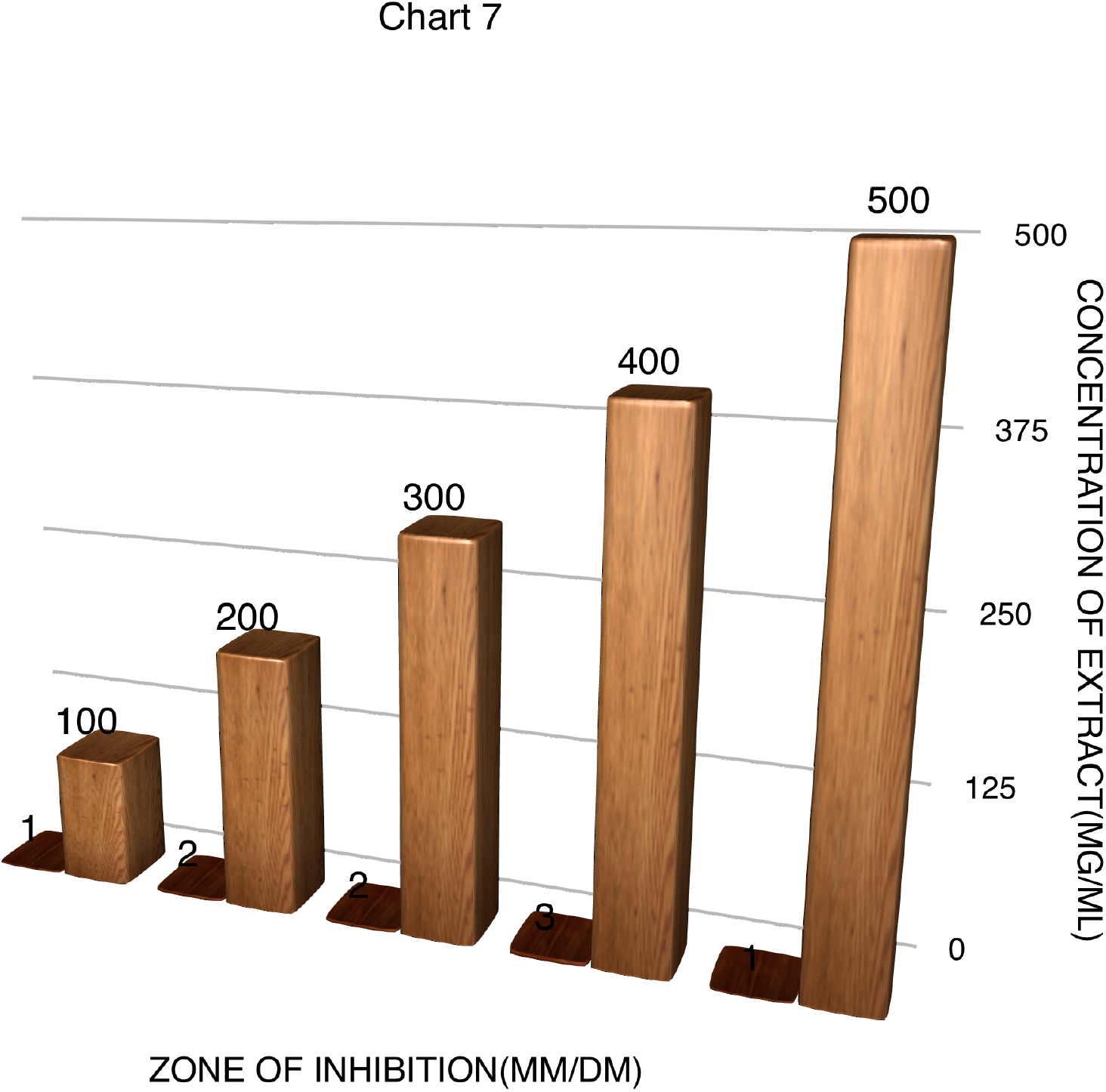

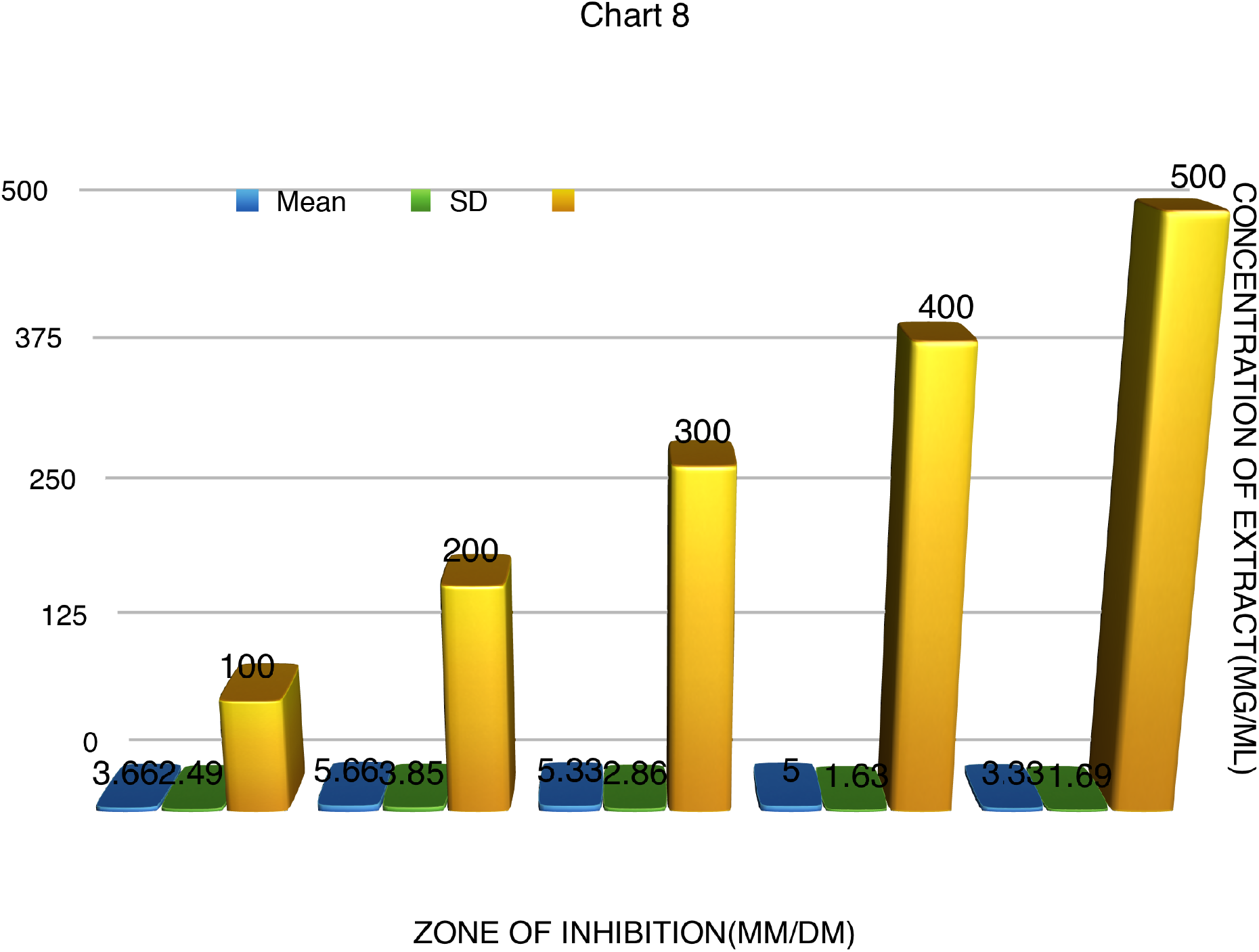

## REFERENCE

Bello, O.M., Zaki, A.A., Khan, I.S., Fasinu, P.S., Ali, Z., Khan, I.A., Usman, L.A., Oguntoye, O.S., (2017). Assessment of selected medicinal plants indigenous plants indigenous to West Africa for antiprotozoal activity. S. Afr. J. Bot. http://doi.org/10.1016/j.sajb.2017.08.002.

Brent, A.J., Oundo, J.O., Mwangi, I., Ochola, L., Lowe, B., Berkley, J.A., (2006). Salmonella bacteremia in Kenyan children. Pediatr. Infec. Dis. J. 25, 230–236.

Chaturvedi, H.K., Mahanta, J., Pandey, A., (2009). Treatment-seeking for febrile illness in north-east India: an epidemiological study in the malaria endemic zone. Malaria J.. 8, 301.

Church, J., Maitland, K., (2014). Invasive bacterial co-infection in African children with Plasmodium falciparum malaria: a systematic review. BMC Med. 12, 11–16.

Croft, S.L., Yardley, V., (2002). Chemotherapy of leishmaniasis. Curr. Pharm. Des. 8, 319–342.

Daya, L., Chothani, H., Vaghasiya, U., (2011). A review on *Balanites aegyptica* Del (desert date): phytochemical constituents, traditional uses, and pharmacological activity, Pharmacogn Rev. 5 (9), 55–62.

Doughari, J.M., Pukuma, M.S., De, N., (2007). Antibacterial effects of *Balanites aegyptica* L, Del, and *Moringa oleifera* Lam, on *Salmonella typhi*. Afr J. Biotechnol.. 6, 2212–2215.

Evans, J.A., Adusei, A., Timmann, C., May, J., Mack, D., Agbenyega, T., Horstmann, T., Frimpong, E.R.D., (2004). High mortality of infant bacteraemia clinically indistinguishable from severe malaria. QJM - Int. J. Med. 97, 591–597.

Gwer, S., Newton, C.R., Berkley, J.A., (2007). Over-diagnosis and co-morbidity of severe malaria in African children: a guide for clinicians. Am. J. Trop. Med. Hyg. 77, 6–13.

Karuppusamy, S., Rajasekaran, K.M., Karmegam, N., (2002). Antimicrobial activity of *Balanites aegyptiaca* (L.) Del. J. Ecotoxicol. Environ. Monit. 12 (1), 67–68.

Natarajan, Devarajan, Ramachandran, Andimuthu, Srinivasan, Kesavan, Mohanasundari, Chokkalingam, (2010). Screening for antibacterial, phytochemical and pharmacognostical properties of *Indigofera caerulea* Roxb. J. Med. Plants Res. 4 (15), 1561–1565.

Noor Jahan, Mohd. Shahid, Shahzad, Anwar, Sahai, Aastha, Sharma, Shivali, Parveen, Shahina (2012). Antimicrobial Potential of *Balanites aegyptiaca* (L.) Del, *Stevia Rebaudiana* (Bert.) *Bertoni, Tylophora Indica* (Burm.f.) Merrill, and *Cassia Sophera* (Linn.). Open Conf. Proc. J. 3, 63–69.

Uneke, C.J., (2008). Concurrent malaria and typhoid fever in the tropics: the diagnostic challenges and public health implications. J. Vector Borne Dis. 45, 133–142.

